# Phenotype to genotype mapping using supervised and unsupervised learning

**DOI:** 10.1101/2022.03.17.484826

**Authors:** Vito Paolo Pastore, Ashwini Oke, Sara Capponi, Daniel Elnatan, Jennifer Fung, Simone Bianco

## Abstract

The relationship between the genotype, the genetic instructions encoded into a genome, and phenotype, the macroscopic realization of such instructions, remains mostly uncharted. In addition, tools able to uncover the connection between the phenotype with a specific set of responsible genes are still under definition. In this work, we focus on yeast organelles called vacuoles, which are cell membrane compartments that vary size and shape in response to various stimuli, and we develop a framework relating changes of cellular morphology to genetic modification. The method is a combination of convolutional neural network (CNN) and an unsupervised learning pipeline, which employs a deep-learning based segmentation, classification, and anomaly detection algorithm. From the live 3D fluorescence vacuole images, we observe that different genetic mutations generate distinct vacuole phenotypes and that the same mutation might correspond to more than one vacuole morphology. We trained a Unet architecture to segment our cellular images and obtain precise, quantitative information in 2D depth-encoded images. We then used an unsupervised learning approach to cluster the vacuole types and to establish a correlation between genotype and vacuole morphology. Using this procedure, we obtained 4 phenotypic groups. We extracted a set of 131 morphological features from the segmented vacuoles images, reduced to 50 after a tree-based feature selection. We obtained a purity of 85% adopting a Fuzzy K-Means based algorithm on a random subset of 880 images, containing all the detected phenotypic groups. Finally, we trained a CNN on the labels assigned during clustering. The CNN has been used for prediction of a large dataset (6942 images) with high accuracy (80%). Our approach can be applied extensively for live fluorescence image analysis and most importantly can unveil the basic principles relating genotype to vacuole phenotype in yeast cell, which can be thought as a first step for inferring cell designing principles to generate organelles with a specific, desired morphology.

## Introduction

Cellular organelles like nucleus, mitochondria, and vacuoles are compartments specialized for performing biochemical reactions and their shape and structure relate to the biological functions they mediate. Therefore, being able to design a particular organelle morphology would mean to govern and influence the outcome of certain biochemical reactions [1]. Indeed, engineering cells for producing chemical compounds outside their evolutionary achieved function or for specific medical applications is already an expanding field. Examples are the artemisin antimalaria drug production, which is cheaper and faster when engineered yeast strains are used [2], or the chimeric antigen receptor (CAR)-T cells employed in cancer therapy [3]. However, the mechanistic realization of specific *ad hoc* cellular and organelle morphology is still elusive.

Recent computational advances have produced important novel theoretical paradigms for the computational design of cellular structures [4]. A combination of forward and data-driven modeling represents a possible path towards establishing rational design principles of cellular structures. While much work has been devoted to using mathematical models to investigate design principles of living organisms, generally considered part of Systems Biology, the same is not true for a purely data driven approach. Specifically, data driven design of living organisms must be achieved through a precise characterization of the relationship between genetic modifications and the resulting changes in morphology. This is a complicated problem of multiplicity: many distinct morphologies can be associated to more than one single driving mutation. Thus, a map of the relationships linking desirable structures to a set of mutations capable of generating those structures is a first, important, and necessary stepping stone towards more complex problems, like the effect of combinations of mutations or environmental perturbations.

A systematic mapping of cellular morphology to mutations in model organisms like yeast is not itself a new problem. Recent papers have started to generate these approaches for several organelles using various supervised unsupervised machine learning techniques [5–8]. However, because of sheer biological variability, the establishment of a large and comprehensive training set (annotated or not) is evidently an important bottleneck. In order to overcome this issue, we introduce a hybrid supervised-unsupervised learning approach which aims at performing accurate end to end mapping between established morphologies and genetic insults in budding yeast organelles with minimal supervision using a small initial training set. We use the yeast vacuole as a test organelle.

Our methodology, which is composed of several steps, is described in Fig. 1 and can be summarized as follows: after the cellular images are acquired, a segmentation step is performed using a U-Net CNN trained on a small fraction of the dataset. The segmented images are then analyzed and a set of features is extracted from the masks. Images are partitioned in an unsupervised fashion with the number of classes automatically inferred from the data, and the resulting clusters are used as training set for another CNN, which is ultimately used to classify the cells. Our results suggest that this methodology can reveal the relationship between phenotype and genotype in an accurate and unbiased way. In the next Section we describe the data acquisition procedure and methods.

**Figure 1:**
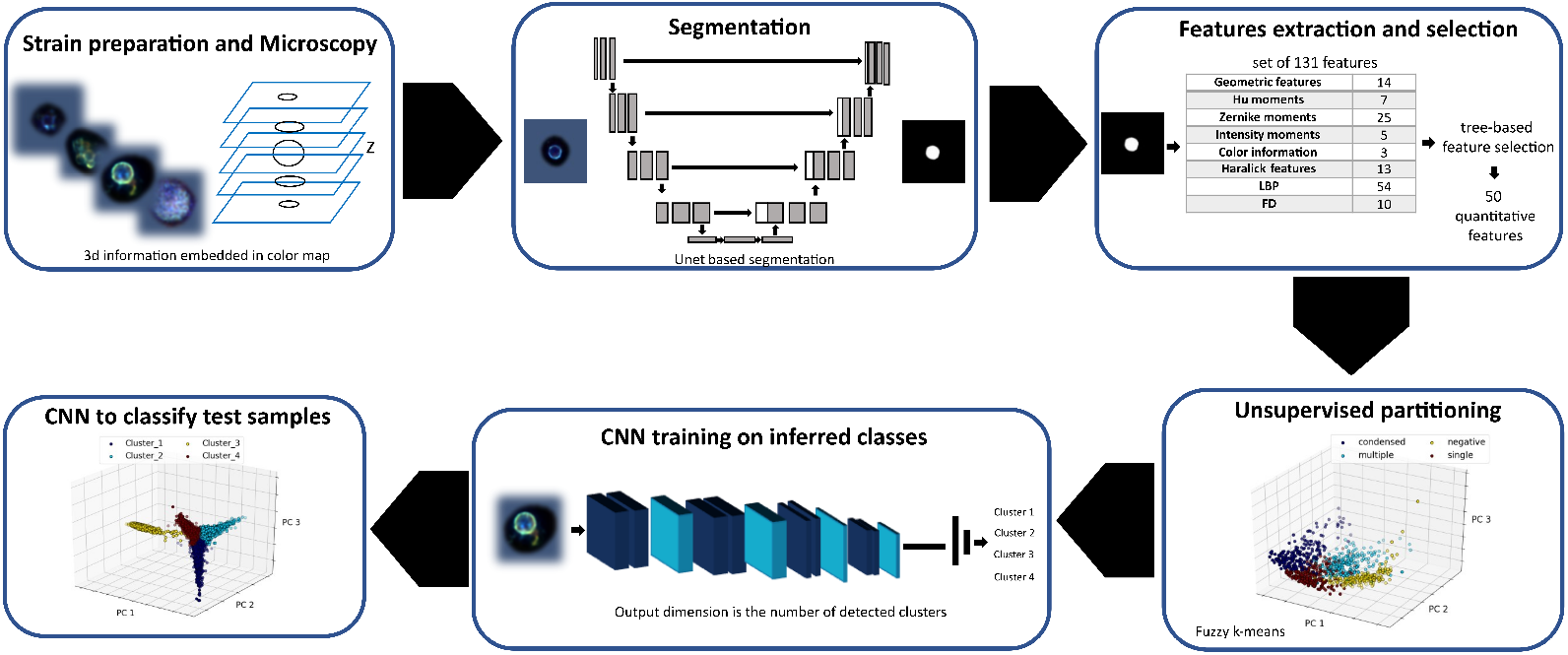
Schematic description of our end-to-end pipeline.

## Materials and Methods

### Dataset description

We used the yeast strain BY4741 VPH1-GFP-HIS3 [9]. Prior to imaging, 96 well glass bottom plates (Cat# MGB096-1-2-LG-L, Dot Scientific Inc.) were coated with 0.1 mg/mL Concavalin A (Sigma-Aldrich, St. Louis, MO) to allow live yeast cells to stick to the bottom. After growing overnight to saturation, cells were diluted and grown to OD 0.4 in YPD media supplemented with 5 mM adenine and 10 mM uracil, then spun at 2000 x g for 2 minutes. The media was removed and cells were resuspended in PBS buffer. The cells were added carefully to the wells of the glass bottom plate and allowed to settle for 2 minutes. Next, the unbound cells were carefully pipetted out along the corner of the wells and fresh PBS was added to the wells. The plates were imaged using a GE InCell 6000 imaging platform (Cytiva, Marlborough, MA). A 2D brightfield image was taken to use for yeast segmentation (100 ms exposure). Forty-eight 0.2 *μ m* sections of VPH1-GFP fluorescence were used to obtain 3D vacuole structure (60X magnification, N.A. 0.95 at 100 *ms* exposure). In order to mitigate distortion of the cell images due to a refractive index mismatch between the air objective lens and the aqueous sample, images were deconvolved with calculated point spread functions that account for this mismatch. Deconvolution was done using a maximum entropy algorithm [10]. A collection of 96 yeast knockouts, *i. e*. complete suppression of the genes, transformed with VPH1-GFP were imaged for a total of 8022 vacuole single cell images, which were used for our study. Each image is 80 × 80 pixels in size and contains a single cell. The image data was parsed from the InCell XDCE metadata file using an XDCE parser module written in Python. A U-net was trained to segment yeast cells in brightfield in Keras (via Tensorflow backend) [11]. Using this model, the predicted 2D masks were generated for each image and a cookie-cutter module was used to ‘ cut’ out individual cells from the field. This information was used to reconstruct the 3D volume. In order to retain the 3D information within a 2D image, a depth-dependent color encoding was performed. This was done by normalizing the image intensity from 0 to 1 and assigning a unique color to each z slice. Therefore, the vacuoles resulted to be colored to encode for depth, blue being at the bottom and red being at the top (Fig. 4).

## Results

### Image segmentation

The first step of the proposed pipeline consisted in a deep-learning based vacuoles segmentation. We used ImageJ [12] to annotate a subset of 500 images randomly extracted from the complete set of vacuole images. We trained an ImageNet pre-trained customized version of the U-Net architecture [11] on the annotated data. Finally, we used the trained model for the segmentation of a data set composed of 1080 yeast vacuoles images, which we used to test our pipeline.

### Features extraction and selection

The output of the segmentation network is an image where only vacuoles are present. Starting from such masked images, we engineered a set of 131 morphological features to describe the shape and texture of the yeast vacuoles. The designed set included: 14 geometric features (e. g. area, perimeter, circularity and eccentricity) 7 moments-based features (using Hu-moments [13]) and 25 Zernike moments (up to order 5 [14]). In addition, we extracted 10 Fourier Descriptors (FD) from the vacuoles contour to further infer the diverse vacuole morphology. To compute the FD, we processed the contour using a shape signature consisting in subtracting the correspondent centroid. To distinguish the yeast vacuole graylevel patterns, we implemented a set of texture-based features, which include 54 Local Binary Pattern (LBP) and 13 Haralick descriptors [15]. A set of 5 features is extracted from the gray-values histogram (mean value, standard deviation, kurtosis, skewness, and entropy) [16]. Finally, we used the ratios between the mean pixel intensity values of the three color channels as features to complete the set of 131 engineered descriptors. Because color includes information on the 3D spatial distribution of the organelles in our 2D image and is encoded in the image texture, the texture-based features are particularly relevant (see Fig. 2. To select only the most important features and avoid performance degradation due to correlation, we adopted a decision tree-based approach using a sci-kit learn implementation [17]. Fig. 2 shows the computed feature importance score. The feature selection procedure results in a final set of 50 features, both shape- and texture-based. Notably, the FD are automatically discarded because of their low score.

**Figure 2:**
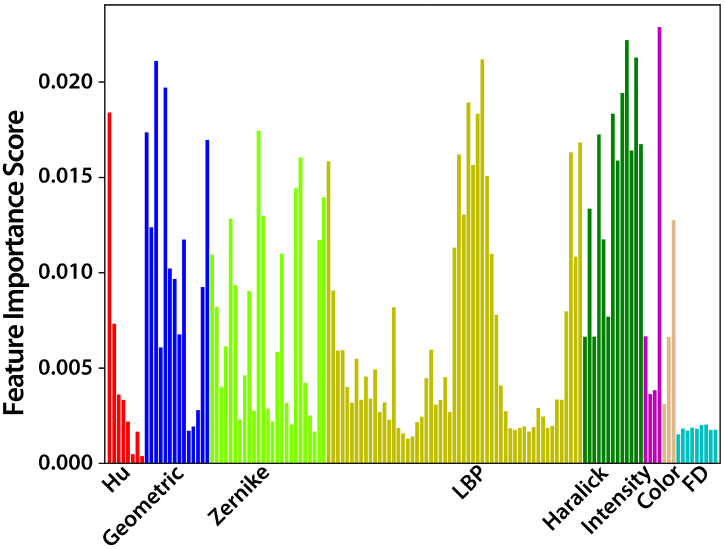
Feature importance computed using a decision tree-based approach on the training set.

### Unsupervised partitioning

From the entire set of vacuole images, we used a subset of 1080 images for the partitioning module of our pipeline and split this subset in an 80:20 ratio for training and testing. We used the features extracted from the training set as input for a Fuzzy K-Means clustering algorithm [18], chosen because of the fuzziness of the acquired images. We then employed a partition entropy (PE) algorithm to infer automatically the number of different morphological classes from the data. The PE coefficient is defined as:

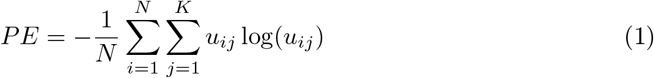

where *u*_*ij*_ is the degree of membership of the sample *i* to the cluster *j, N* is the total number of samples, and *K* the maximum number of considered clusters. The PE coefficient varies between 0 and log(*K*), with higher uncertainty in clustering for lower PE values. Therefore, a good estimation of the number of classes corresponds to the PE peak with respect to *j*.

We evaluated the accuracy of the clustering algorithm by means of a purity metric [19], defined as:

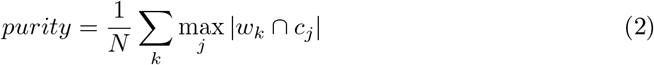

and assigned the ground truth class *k* to the most overlapping cluster *j*.

### Evaluation of the testing image subset using the implemented CNN and deep feature extraction

From the unsupervised partitioning we obtained a set of labels, which assign each of the training sample *x*_*i*_ to one cluster *y*_*j*_. We used this annotated set to train a CNN using a Stochastic Gradient Descent (SGD) optimizer. We then used the trained model to evaluate the 200 image testing set. We performed deep feature extraction by adding a layer of 40 neurons (dashed layer in Fig 3). In this way we build a feature space that corresponds to the class boundaries learnt by the trained model.

**Figure 3:**
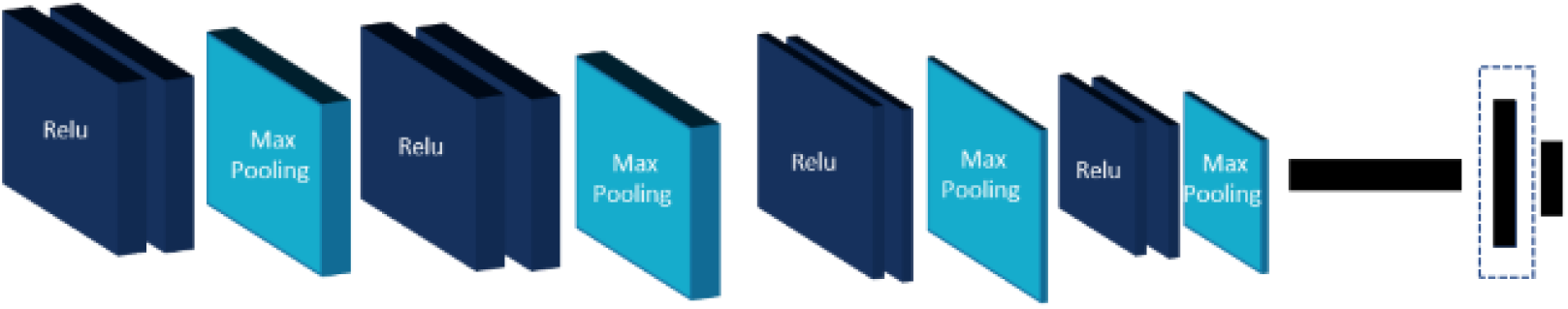
Representation of the CNN architecture. The dashed rectangular shape indicates a 40 neuron layer used for deep feature extraction.

## Results

### 0.1 High accuracy of the training set clustering procedure

Using PE we find that 4 classes best divide the training data set. We then used the Fuzzy K-Means algorithm to cluster this data set and evaluate the purity of the clustering procedure using our team’ s expert annotation. The Fuzzy K-Means algorithm showed a purity of ≈ 85% and the 4 detected clusters correspond to 4 specific morphological classes (see Fig. 4): single vacuole, multiple (2 or more) vacuoles, condensed vacuoles, and dead cell, which we call negative. To provide a graphical representation of the algorithm accuracy, we performed a Principal Components Analysis (PCA) on the Fuzzy K-Means input features. Figs 5A-B show the space of the first 3 principal components using the ground truth (Fig 5A) and the annotated labels (Fig 5B). The cluster colors are assigned evaluating the most overlapping ground truth labels for each of the detected clusters. The two graphs are in excellent agreement, proving high accuracy of the unsupervised partitioning procedure. To evaluate the effect of feature selection on the clustering accuracy, we tested the Fuzzy K-Means algorithm with both the 131-dimensional feature set (the total set of hand-engineered features) and the 50-dimensional set resulting from the decision-tree based feature selection on the total set, and we obtained a maximum value of 82% and 85%, respectively.

**Figure 4:**
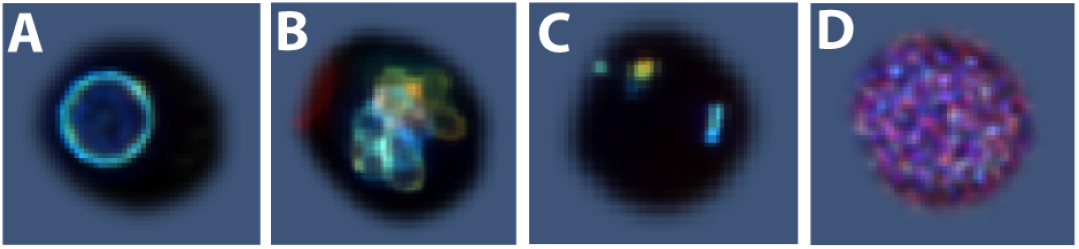
Representative image for the 4 classes identified by the PE: A) single, B) multiple, C) condensed, and D) negative

**Figure 5:**
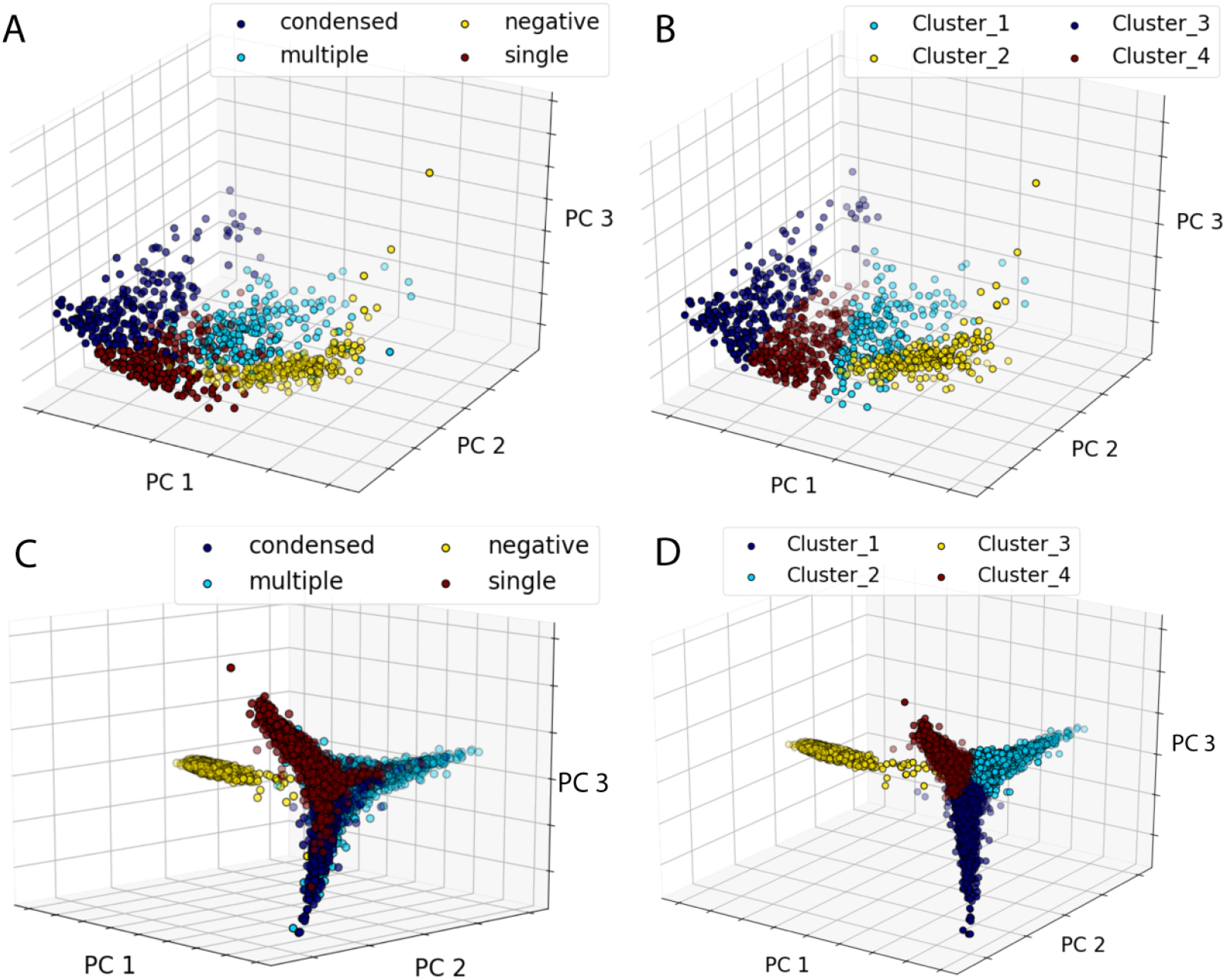
Comparison of the principal component (PC) analysis of the data set used for the unsupervised partitioning calculated with the classes annotated by our team’ s expert A) and assigned by the clustering algorithm B). Comparison of the PC analysis of the deep features extracted using the CNN with classes annotated by our team’ s expert C) and determined by the clustering algorithm D).

### High accuracy of our CNN model

The trained CNN reaches a validation accuracy of 99% and a testing accuracy of ≈ 81%. We then used our CNN to predict a larger non-annotated testing set containing 6942 images. The model partitioned the test set into the four morphological classes with the following occurrences: 1618 single vacuole, 2727 multiple, 1477 condensed, and 1120 negative, resulting in a strongly imbalanced set. To estimate the accuracy of the trained model, we asked an expert to review the classification results in terms of the 4 classes of vacuoles previously described, computing the consequent accuracy. The trained model used on the whole set showed an accuracy similar to that of the smaller testing set (80%), suggesting that the implemented pipeline can be used with high confidence to test significantly larger data sets without annotation. This result is remarkable because the annotation of biological images generally suffers from annotation bias, *i*.*e*., there is inherent variability due to annotation of different experts. Our pipeline might help mitigate this problem. For instance, in our specific case, the 20% of false positives are mostly multiple vacuoles classified by our team’ s expert as condensed (645 cells) or single (297 cells). To provide a visualization of the class boundaries learnt by the trained CNN, we performed a PCA on the deep extracted features and we represented the results in Figs. 5C and 5D. Fig. 5C shows the ground truth labels and Fig 5D the labels assigned by the CNN prediction. The two graphs are very similar, as expected considering the high CNN accuracy. We built the ground truth of Fig. 5C considering that the CNN provided true positives and our team’ s expert re-labeled false positives. As a benchmark, we trained a CNN directly on the ground truth labels obtaining a testing accuracy of 85%, very close to the accuracy obtained with our unsupervised approach.

### Relating vacuole morphology to different genetic conditions

In Fig. 6 we report the relative number of cells belonging to each class as a function of the 96 wells (see Dataset description). Each panel represents a class and the elements of the matrix correspond to the experimental wells and thus to specific mutations. The color represents the relative fraction of a particular morphological type. Our results show that different genetic perturbations result in similar morphological types with a measurable frequency. This shows the potential for our pipeline to suggest several mutations capable of achieving a specific phenotype.

**Figure 6:**
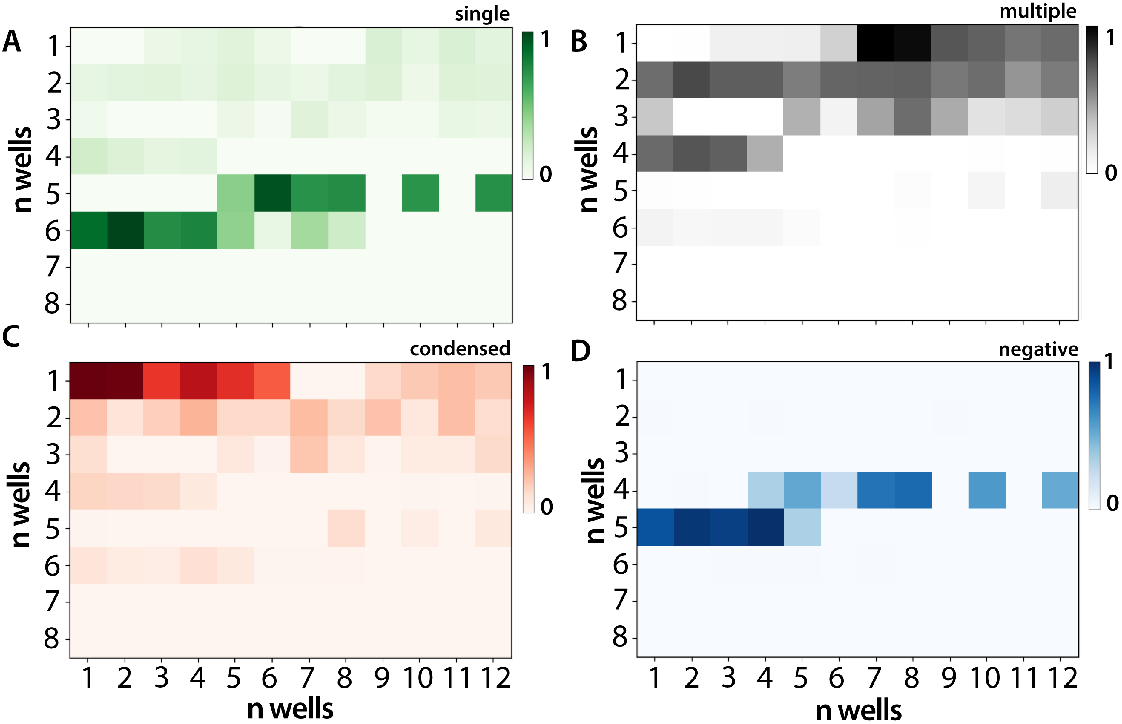
Normalized frequency of occurrence of A) single vacuole, B) multiple, C) condensed, and D) negative as a function of the 96 wells, each of them corresponding to a different phenotype.

## Conclusions

While the mechanistic design of cells and cellular structures is still a long term scientific goal, introducing methods capable of inferring biological design principles is an important first step towards mechanistic realization of biological systems for useful purposes. The yeast vacuole is an example of an organelle which can be used as a biological reactor for chemical production. Our pipeline is capable of relating distinct vacuole morphologies to genetic perturbations. For this purpose we adopt a mixed supervised-unsupervised learning methodology with the aim of reducing the annotation burden and the inherent bias due to the human annotation task. Our results show that the link between cellular phenotype and genotype can be obtained with high accuracy with limited user intervention. While the investigation of more complex engineering tasks, like creating desired cellular structures using multiple co-occurring mutations or modification of the cellular environment, has not been addressed in this work, we believe our investigation provides the necessary background for a more general understanding of the engineering process.

### Broader Impact

The investigation of the relationship between genotype and phenotype with the aim of establishing design principles of rational engineering of cells and cellular structures is an important aim of today’ s biotechnology. While on one hand it is important to perfect the experimental realization, it is paramount to achieve precise control of the design task. The routine trial and error cycle of synthetic engineering is expensive, inaccurate and incremental. Our broader aim is to understand how machines can help not only accelerate design, but rather and more importantly enable the mechanistic realization of useful biological products. From the point of view of the societal impact, this larger scope, of which our methodology represent a first fundamental step, may open a whole new industry for which a new workforce needs to be trained. As more and more computational methods become part of the bioengineering playbook, new professional figures will emerge, which will benefit the society as a whole. From an ethical perspective, the importance of achieving desired products and exploring a larger space of possible outcomes reduces the uncertainty of the consequences of the synthetic engineering process. While this uncertainty cannot be completely removed from the equation, a more controlled design is a desirable feature of any pipeline which aims at building living matter.

## Acknowledgments

We thank Brenda Andrews for sending us her list of gene knockouts that resulted in changes in vacuole morphology. This material is based upon work supported by the National Science Foundation under Grant No. DBI-1548297

